# In situ genotyping of a pooled strain library after characterizing complex phenotypes

**DOI:** 10.1101/142729

**Authors:** Michael J. Lawson, Daniel Camsund, Jimmy Larsson, Özden Baltekin, David Fange, Johan Elf

## Abstract

So far, it has not been possible to perform advanced microscopy on pool generated strain libraries and at the same time know each strain’s genotype. We have overcome this barrier by identifying the genotypes for individual cells *in situ* after a detailed characterization of the phenotype. The principle is demonstrated by single molecule fluorescence imaging of *E. coli* strains harboring barcoded plasmids that express a sgRNA which suppress different genes through dCas9.

---

Recent years have seen a rapid development in genome engineering, which, in combination with decreased costs for DNA oligonucleotide synthesis, have made it possible to design and produce pool generated cell libraries with overwhelming genetic diversity^1–6^. A similarly impressive development in microscopy enables the investigation of complex phenotypes at high temporal resolution and spatial precision in living cells^7,8^. Finally, developments in microfluidics have enabled well controlled single cell observations of individual strains over many generations^9^. Despite the rapid technological progress within these areas, there is currently no efficient technique for sensitive time-resolved phenotyping of pool generated libraries of genetically different cell strains. Recent work observing multiple bacterial strains on agarose pads allows for sensitive microscopy^10,11^, but the genetic diversity is capped since the strain production and handling is not pooled. On the other end, droplet fluidics allow working with large genetic diversity^1^, but cannot be used to characterize phenotypes that require sensitive time lapse imaging.

Our solution is composed of three key components: (1) construction of a barcoded strain library, (2) live cell phenotyping in a microfluidic device where each strain occupies a defined position and (3) *in situ* genotyping to identify which strain is in which position (Fig. 1). Phenotyping is here used as a general description for any of the time-resolved biochemical assays that are possible with time-lapse microscopy in individual cells. We refer to the method as DuMPLING - Dynamic u-fluidic Microscopy based Phenotyping of a Library before IN situ Genotyping.

**Figure 1.**
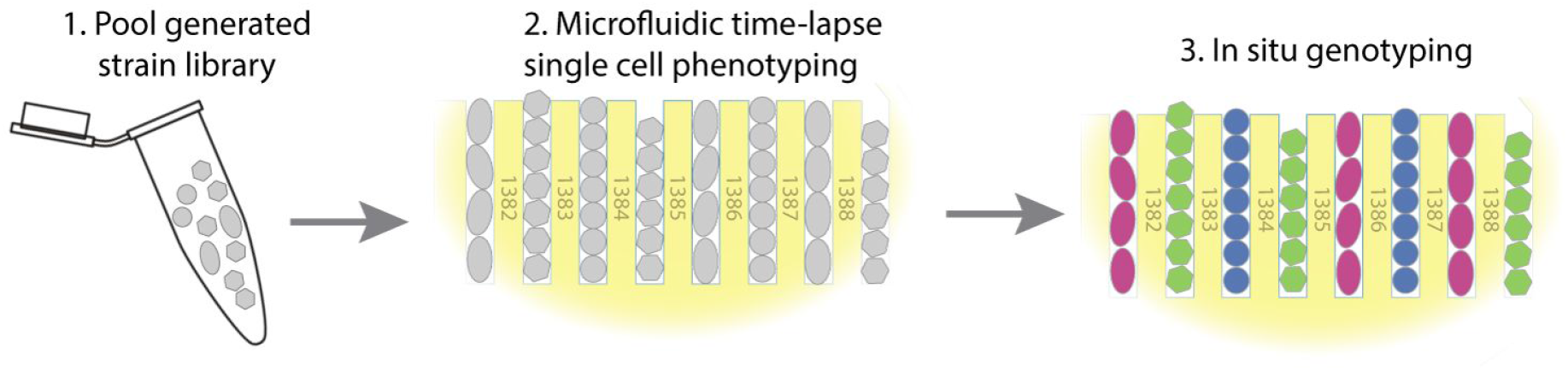
The DuMPLING strategy: (1) pooled strain library generation of e.g. dCas9 knockdowns or chromosomal promoter variants. (2) Live cell phenotyping using fluorescence microscopy to e.g. identify changes in protein localization, diffusion, or kinetics. (3) Genotypes recovered by *in situ* genotyping using e.g. FISH or *in situ* sequencing.

Each component has previously been successfully developed and can be performed in many different ways. For example, one can make pooled dCas9 libraries based on plasmids harboring both a genotype-identifying barcode and a sgRNA gene^1–5^ for labeling genetic loci^12^ or knocking down/activating genes throughout the chromosome. Alternatively, pooled chromosomal libraries with variants of promoters, ribosome binding sites or coding sequences^6,13^ can be made. Similarly, single cell time courses have been obtained with high sensitivity in dedicated microfluidic devices^8,14–17^. The method for identifying the barcode can be based on *in situ* sequencing^18,19^ or sequential probing using fluorescence *in situ* hybridization (FISH)^20,21^.

To exemplify the use of DuMPLING, we targeted different components of the *lac* operon in *E. coli* using a set of sgRNA-expressing plasmids that repressed *lacI, lacY* or an unrelated gene (Fig. 2A). The plasmids are made from pooled oligos including the sgRNA and its unique barcode. The pooled approach has previously been used to generate libraries of thousands of genotypes^1,3^, but here we limit to three variants to be able to precisely evaluate the accuracy of each step. The mixed plasmids are electroporated into an *E. coli* strain, where dCas9 is expressed from a regulated chromosomal promoter. Furthermore, the *lacY* gene is fused with the gene for the fluorescent protein YPet to obtain an easily detectable phenotype.

The mixed strains are loaded into a microfluidic chip which harbours 2000 cell channels, sustains continuous exponential growth, and allows single cell imaging for days (Fig 2B). After a few generations, all cells in a channel are the progeny of the cell at the back of the channel and thus share the same genotype. The chip design is similar to the mother machine^9^, but we have introduced a 300 nm opening in the back of each cell channel such that media and reagents can be passed over the cells. In our experiments, 233 channels are imaged every 60 seconds using phase contrast and every 13 minutes using single molecule sensitive wide field fluorescence for a total of 272 minutes. Phase contrast images are used for cell detection and lineage tracking. Individual LacY-YPet molecules, detected using wide field epifluorescence, are overlaid on the phase contrast images to allow assignment of individual molecules to individual cells. At this point the phenotypes are only associated to the spatial position in the chip (i.e., the channel the cells are in), since the genotype of the cells is not yet known.

Each plasmid expresses a unique RNA-based barcode that allows genotype identification. The barcode is expressed from a T7 promoter, and the T7 polymerase is under control of an inducible arabinose promoter. The orthogonal and inducible nature of this system prevents it from interfering with cell growth before induction (Fig. S3). The cells are fixed *in situ* with formaldehyde and permeabilized before FISH. The individual barcodes are identified by sequential hybridization of fluorescent 37 base long oligonucleotides (probes). The multiplexed process of designing and producing the probe library is described in the Supplementary Methods. We use probes of two different colors (C=2) in two sequential rounds of probing (N=2), which is sufficient for identifying the 3 genotypes in this study. In general, C^N^ genotypes can be identified. To demonstrate that it is possible to reprobe many times, we perform N=6 consecutive rounds (Fig. 2C) of probing in each position. Extending to C=4 colors without significant bleedthrough based on currently available dyes is common practice^19^, and therefore it is possible, without significant additional engineering, to uniquely encode and identify >4000 strains.

**Figure 2.**
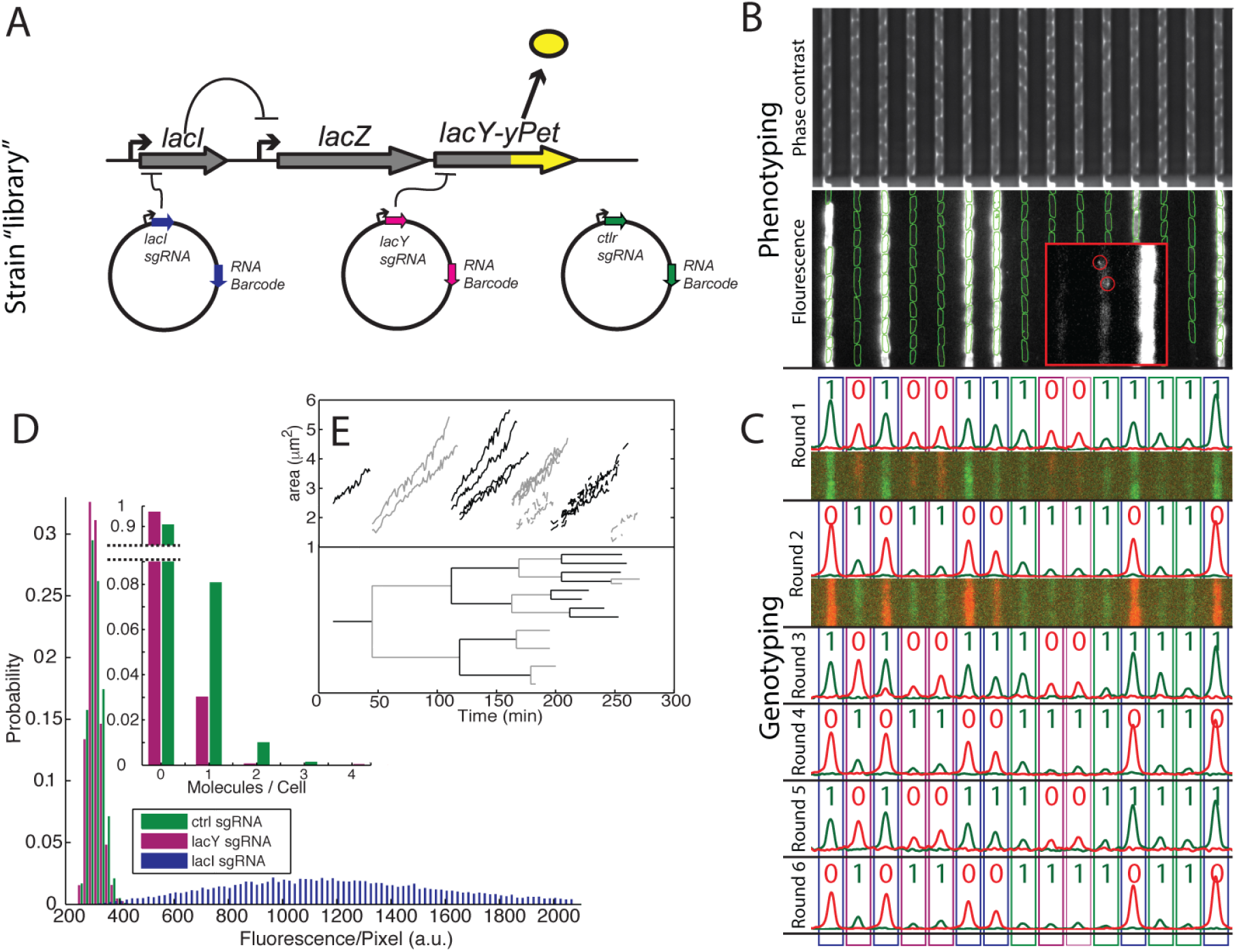
Example experiment. **A**. Repression network for for the three different plasmids used. **B**. Examples of channels and cells in the custom-made microfluidic device which are imaged in both phase contrast and fluorescence microscopy. Phase contrast is used to segment the cells and single molecule fluorescence microscopy is used to detect gene expression from the *lac* operon. **C**. *In situ* genotyping with 6 sequential rounds of FISH probe hybridization. **D**. Gene expression categorized by assigned genotype. *Inset* Single molecule counting of expression from the two low expression genotypes. **E**. *Top* growth curves for one cell lineage (from one channel). Dashed lines indicate the end of detection of a branch. *Bottom* corresponding lineage tree.

In Fig. 2D we show the distribution of the number of LacY-YPet detected per cell in individual frames for the different genotypes. As a control for correct genotype to phenotype assignments we note that all strains that express a high level of LacY-YPet have been correctly found to express the barcode RNA associated with the sgRNA against *lacI* and that all strains with the barcode RNA associated with the sgRNA against *lacI* express high levels of LacY-YPet. As a demonstration of single molecule sensitivity in the phenotyping, we note that we can reproducibly measure mean expression of less than one YPet molecule per generation and distinguish a less than 3x change at this expression level (2D *inset*, Fig. S6). We also demonstrate the dynamic capability by plotting an example cell lineage tree that spans 6 generations and the full time course of the experiment (Fig. 2E).

This brief communication describes a proof of principle application of the DuMPLING concept, i.e. the possibility to phenotype a pool generated library of cells and then genotype *in situ*. The advantage of our method compared to the state of the art is the combination of pooled handling of library generation and characterization of complex phenotypes based on dynamic changes in single cells. We have here restricted ourselves to a small CRISPRi application with bacteria in a fluidic device, but the dumpling can have many other fillings.

## Acknowledgements

The authors would like to thank George Church and Mats Nilsson for helpful discussion. In addition, the authors would like to acknowledge funding from the Knut and Alice Wallenberg foundation, the Swedish Research Council and the European Research Council.

